# Production and assessment of monoclonal antibodies against the SpyCas9 protein of *Streptococcus pyogenes*

**DOI:** 10.1101/2021.01.02.425082

**Authors:** Mi-Jin Park, Jongjin Park, Slki Park, Sunghwa Choe

## Abstract

The biotechnological applications of the programmable DNA-cleaving enzymes known as clustered regularly interspaced short palindromic repeats/CRISPR-associated protein 9 (CRISPR/Cas9) continue to expand from agricultural trait development to therapeutic gene therapies. The Cas9 CRISPR system isolated from *Streptococcus pyogenes* (SpyCas9) is classified as class II, which is popular due to high efficiency and robustness. Antibodies that specifically detect SpyCas9 could expand the applicability of this enzyme. Here, we report on the development of monoclonal antibodies against SpyCas9. Four hybridoma cells were selected for their expression of anti-SpyCas9. Hybridoma supernatant contained reactive and highly specific antibodies to SpyCas9. Anti-SpyCas9 antibody was purified and was non-selective against six other bacterial Cas9 proteins. SpyCas9 protein could be detected in HEK293T cells in decreasing amounts over a 48 hour period, indicating the antibody could be used to detect residual levels of SpyCas9 remaining following cell treatment with a CRISPR/SpyCas9 system. The anti-SpyCas9 antibody may help researchers facilitate Cas9 for further study for the use of techniques such as enzyme-linked immunosorbent assays (ELISAs), western blot, immunoprecipitation, and immunohistochemical staining.

## Introduction

Clustered regularly interspaced short palindromic repeats/CRISPR-associated protein 9 (CRISPR/Cas9) were first reported in 2012. CRISPR/Cas9 activity results in double-stranded breaks (DSBs) in DNA by binding to a specific target site, which is decided by a protospacer that has an antisense sequence for the specific target site (Burley and Sedgley, 2012). Foreign DNA cleavage by CRISPR/Cas9 in bacteria is used as defense system against viral or bacteriophage infection (Datsenko et al., 2012), producing a sophisticated immune system. The cleavage function of CRISPR/Cas9 has subsequently been applied for gene-cloning at the laboratory bench (Pickar-Oliver and Gersbach, 2019). CRISPR techniques can disrupt double-stranded DNA to produce knockdown of targeted gene expression as well as induce cell death (Adli, 2018).

To substitute a nucleotide base on the target base without DSB, the base-editor (BE) and prime-editor (PE) versions of this system have been developed. These both utilize nickase Cas (nCas) proteins that are dedicated to bind to the target DNA region with single-guide RNA (Hampton, 2020; Rees and Liu, 2018). The BE uses a cytidine deaminase or an adenine deaminase, which can substitute C to T and an A:T base pair to a G:C base pair when applied to human or plant cells (Eid et al., 2018; Rees and Liu, 2018). The PE employs a reverse transcriptase that replaces X to Y in yeast and rice cells (Anzalone et al., 2019; Tang et al., 2020). In genome editing, quantifiable precise control is required to avoid off-target effects. However, even when non-cleavage editing technologies have been applied, uncontrollable excess usage of CRISPR cleavage using plasmid DNA, mRNA, and protein has generated off-target effects (Zuo et al., 2019).

Control of CRISPR cleavage in cells is a crucial factor to reduce off-targeting. Precise observation for CRISPR cleavage in terms of the balance between genome editing efficiency and off-targeting effects is required. The expression of the CRISPR-associated protein 9 (Cas9) should be precisely detected by a highly qualified anti-Cas antibody independently of which particular CRISPR molecular types are used. An anti-Cas antibody is expected be able to bind to the protein expressed from Cas9 DNA and mRNA. To facilitate the functional study of Cas9 for further characterization when using enzyme-linked immunosorbent assays (ELISAs), western blot, immunoprecipitation (IP), or immunohistochemical staining (IHC), we developed highly qualified anti-*Streptococcus pyogenes* (Spy)Cas9 monoclonal antibodies (MAbs) by using a recombinant Cas9 protein.

## Materials and methods

### Production of recombinant *SpyCas9* protein

The *SpyCas9* gene with a 6×His tags at the N- and C-termini was cloned into the pET28a vector. For bacterial expression of SpyCas9, the expression vector was transformed into Rosetta 2 (DE3) pLysS *Escherichia coli* (Novagen, Madison, WI, USA). An *E. coli* colony harboring SpyCas9 was inoculated in Luria Bertani (LB) medium with kanamycin and cultured at 37 °C overnight. Cultures were then sub-cultured in LB medium and incubated at 37 °C until the optical density at 600 nm (OD600) reached 0.4–0.7. SpyCas9 expression was induced by addition of 1 M isopropyl-β-D-thiogalactopyranoside (IPTG; final concentration 1mM); incubation was continued for 20 h at 18 °C. The culture was centrifuged at 4000 rpm for 30 min and the cell pellet was resuspended with lysis buffer containing 20 mM Tris-HCl (pH 8.0), 5 mM imidazole, 500 mM NaCl, 1 mM 1,4-dithiothreitol (DTT), and 1 mM phenylmethylsulfonylfluoride (PMSF). Resuspended cells were lysed by a probe sonicator and the lysate was subjected to centrifugation at 15,000 × *g* for 1 h. The supernatant was filtered through a 0.45 μm syringe filter and purified using immobilized metal affinity chromatography via a HisTrapTM HP 5 mL (GE Healthcare, Chicago, IL, USA) and an AKTA FPLC system (GE Healthcare). Recombinant protein was eluted using 20 mM Tris-HCl, pH 8.0, 500 mM NaCl, and 500 mM imidazole in a step gradient from 14% to 100%. The eluate was desalted on a column HiPrepTM 26/10 desalting column (GE Healthcare) and eluted with 20 mM HEPES, 150 mM KCl, 1 mM DTT, 10% v/v, glycerol, pH 7.5. Harvested protein was separated via 10% sodium dodecyl sulfate-polyacrylamide gel electrophoresis (SDS-PAGE) to assess purity. The desalted fraction of SpyCas9 protein was concentrated by using Vivaspin turbo 15 concentrators (50,000 MWCO; Sartorius, Goettingen, Germany). The concentrated protein was quantified by Bradford assay and was kept at −80 °C.

### Generation of four hybridoma clones and IgG purification

A-Frontier Cooperation (Seoul, South Korea) prepared four hybridoma clones from mouse spleens after injecting the in-house purified SpyCas9 protein as an antigen. Following the second boost injection, mouse spleen cells were isolated and fused with myeloma cells (Sp2/0Ag14). Four hybridoma clones (24H5, 9G8, 5G11, and 18A4) that secreted SpyCas9 antibody was obtained via screening.

For IgG purification, culture medium grown with the 18A4 hybridoma was passed through a protein A resin column. After washing the column, an elution buffer was added to obtain a fraction containing the bound antibody. The fraction samples were pooled and concentrated. IgG purified antibody samples were loaded on an SDS-PAGE gel in nonreducing and reducing conditions to confirm purity.

### Cell culture and transfection

HEK293T and HeLa cell lines (from American Type Culture Collection, Manassas, VA, USA) were cultured in Dulbecco’s Modified Eagle Medium (Hyclone Laboratories Inc, Logan, UT, USA) supplemented with 10% heat-inactivated fetal bovine serum (Hyclone Laboratories Inc) and 100U/mL of penicillin-streptomycin (Gibco-Invitrogen, Carlsbad, CA, USA). The cells were transfected at 50%–70% confluency using Viafect™ transfection reagent (Promega, Madison, WI, USA) An optimal condition was adjusted by the transfection efficiency and toxicity assessment. HEK293T cells were treated with a 3:1 Viafect™ reagent and an appropriate DNA ratio. For electroporation, we used the 4D-Nucleofector™ System. HEK293T cells (2 × 10^5^) were resuspended with SF buffer (V4XC-2032, Lonza, Basel, Switzerland) and pulsed with purified SpyCas9 (20 pmol).

### Plasmids

Seven Cas9 expressing plasmids were used in the study (Addgene plasmid # 62988, # 68333, # 68334, # 68335, # 68336, # 68703, and # 68705). Each plasmid was purified via the Exprep™ Plasmid SV, Mini Kit (Geneall, Seoul, South Korea).

### Immunoblotting

Following SDS-PAGE, proteins were transferred to a polyvinylidene difluoride (PVDF) membrane (MilliporeSigma, Burlington, MA, USA). The membranes were blocked with 5% milk powder in phosphate-buffered saline (PBS)-Tween 20 buffer for 1 h at RT. The membranes were incubated with undiluted hybridoma supernatants, anti-HA (Invitrogen, Carlsbad, CA, USA), anti-β-actin (MBL international, Woburn, MA, USA), or IgG purified SpyCas9 antibody overnight at 4°C with agitation, followed by incubation with goat anti-mouse IgG conjugated to horseradish peroxidase (Sigma-Aldrich, St. Louis, MO, USA) diluted 1: 5000 in PBS-Tween 20 buffer. Proteins were detected using ECL solution (Bio-Rad Laboratories, Hercules, CA, USA).

### Sequencing of SpyCas9 antibody

Y Biologics company (Deajeon, South Korea) extracted mRNA from the 18A4 hybridoma clone provided to sequence variable regions of the antibody. The extracted mRNA of 18A4 were used for synthesizing cDNAs, which contained heavy chain (VH) and light chain (LH) sequences, which were employed for cloning into pGEM®-T Vector. At least five colonies are sequenced to identify the dominant sequence from colonies.

## Results

### Purity of recombinant SpyCas9 produced for antibody production

To produce a monoclonal antibody to SpyCas9, we purified recombinant SpyCas9 from *E. coli*. We confirmed the protein purity via SDS-PAGE to assess suitability for specific antibody production. Purified protein was observed at the expected size for SpyCas9 (166 kDa) (Fig. 1A); minor amounts of nonspecific proteins were also observed. The purified SpyCas9 was then used for immunization to produce anti-SpyCas9 IgG antibody.

**Fig. 1.**
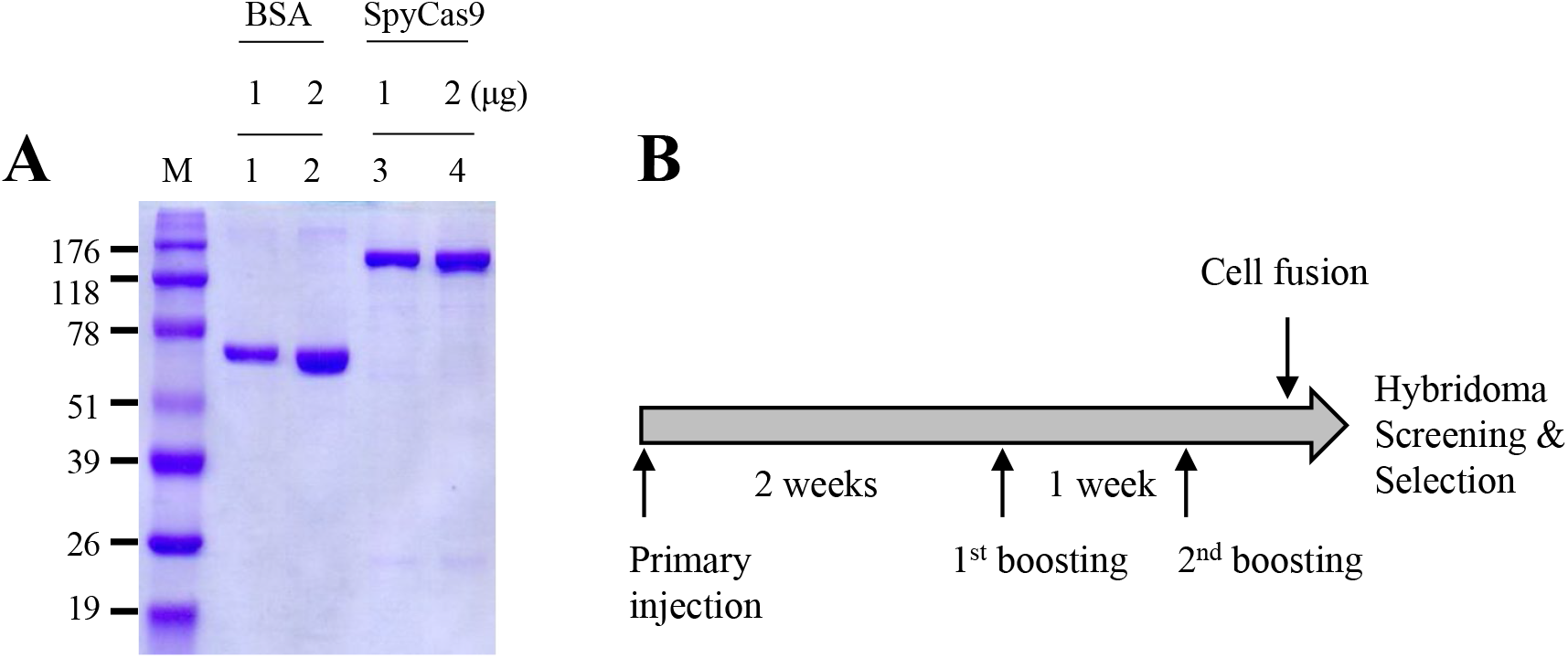
Purification of SpyCas9 and an immunization scheme. (A) Purification of SpyCas9 in *E. coli*. Lane M, protein size standards. Lanes 1-2, a loading control of Bovine Serum Albumin (BSA, 1 and 2 μg); lanes 3-4, purified recombinant SpyCas9 protein (1 and 2μg). The gel was stained with Coomassie Brilliant Blue (CBB). (B) Overview of mouse immunization schedules and hybridoma development protocols for SpyCas9 monoclonal antibody production. Primary and first booster injections are intraperitoneal (IP) as emulsions in Freund’s Complete Adjuvant (CFA). After 2nd boosting, spleen cells are extracted from mouse and fused. Four hybridoma clones are selected.

### Hybridoma cells secreting anti-SpyCas9 IgG antibodies produced via immunization of mice

B cells are well-known antibody-producing cells in vertebrates (Zimmerman et al., 2010). Murine B cells were obtained for producing SpyCas9 antibody by intraperitoneal (IP) injection of recombinant SpyCas9 mixed with Complete Freund’s Adjuvant to female BALB/c mice. In total, three injections were performed for boosting (Fig. 1B). Two weeks after the primary injection, SpyCas9 protein was re-injected to the mouse for the first boost, and the second boost was followed by assessment for the response to immunization. After three days, B cells were isolated from mice spleens, and the isolated B cells were fused to myeloma cells to make hybridoma cells. Hybridoma cells were screened by ELISA analysis and assessed by cloning until the final clone had been verified (Fig. 1B).

### Candidates for anti-SpyCas9 IgG antibody based on four different hybridoma cells could detect SpyCas9 protein

We obtained four different hybridoma cells (named 24H5, 9G8, 5G11, and 18A4) that secreted anti-SpyCas9 antibody. To test the binding affinity of anti-SpyCas9 IgG antibody, these four hybridoma cells were separately cultured in media; culture supernatants that contained anti-SpyCas9 IgG antibodies were then isolated from hybridoma cells by centrifugation. The supernatant was assessed for detection of SpyCas9 at a range of concentrations (12.5, 25, 50, 100, and 200 ng) via western blotting before additional IgG purification; the supernatant from hybridoma 18A4 gave a particularly strong response (Fig. 2A). We then assessed the specificity of anti-SpyCas9 binding to SpyCas9 protein in the presence of nonspecific proteins (0, 5, 30, and 100 μg) from HeLa cell lysate and demonstrated that these proteins do not interfere with the SpyCas9 to the selected antibodies (Fig. 2B).

**Fig. 2.**
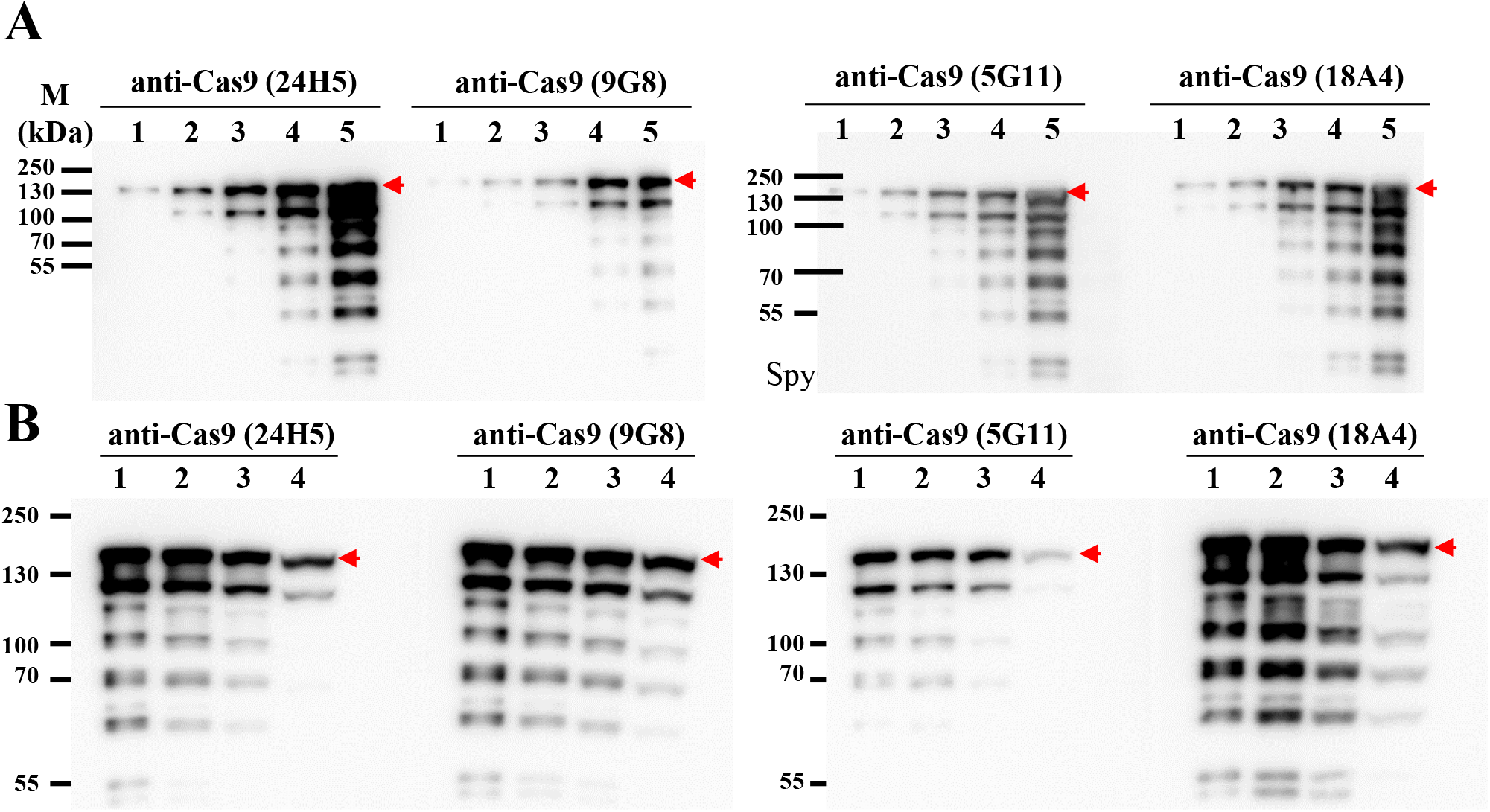
Western blotting; Quantitative and specificity estimation for SpyCas9 antigen. Detection of (A) five different amounts of SpyCas9 (upper panels). Lanes 1-5; SpyCas9 protein of 12.5, 25, 50, 100, and 200 ng. Exposure time for the first two gels and next two gels were 15 and 2 seconds, respectively. (B) A 100 ng purified SpyCas9 without or with nonspecific Hela cell lysate (bottom panels). Each lane was loaded with 100 ng of SpyCas9. Lane 1, without Hela cell lysate; Lanes 2-4, with 5, 30, and 100 μg of Hela cell lysates. Red arrows indicate SpyCas9 protein size and the western blotting was detected by four hybridoma (24H5, 9G8, 5G11, 18A4) culture media. Exposed for 10 seconds.

### Anti-SpyCas9 IgG antibody from hybridoma cell 18A4 detects SpyCas9 protein

From the original four hybridoma cells, we selected hybridoma 18A4 based on detection of SpyCas9 protein by anti-SpyCas9 IgG antibody. The supernatant from hybridoma 18A4 was examined to confirm the binding activity to SpyCas9 protein under different conditions (Fig. 3A). First, crude *E. coli* extracts of SpyCas9 protein were employed (lanes 2–4 of the left and right panels in Fig. 3A) alongside in-house and commercial SpyCas9 proteins (ToolGen, Seoul, South Korea) were tested (lanes 5 and 6 of the left and right panels in Fig. 3A). Recombinant SpyCas9 proteins contained a histidine tag at the C-terminus, enabling detection by of SpyCas9 with anti-His antibody to compare with the results of anti-SpyCas9 detection. Then, the anti-His antibody was stripped off of western blots and the supernatant from the hybridoma cell 18A4 was used for blotting. This confirmed that the supernatant from the hybridoma cell 18A4 detected SpyCas9 protein in spite of potential interference from the *E. coli* lysates (the right panel of Fig. 3A).

**Fig. 3.**
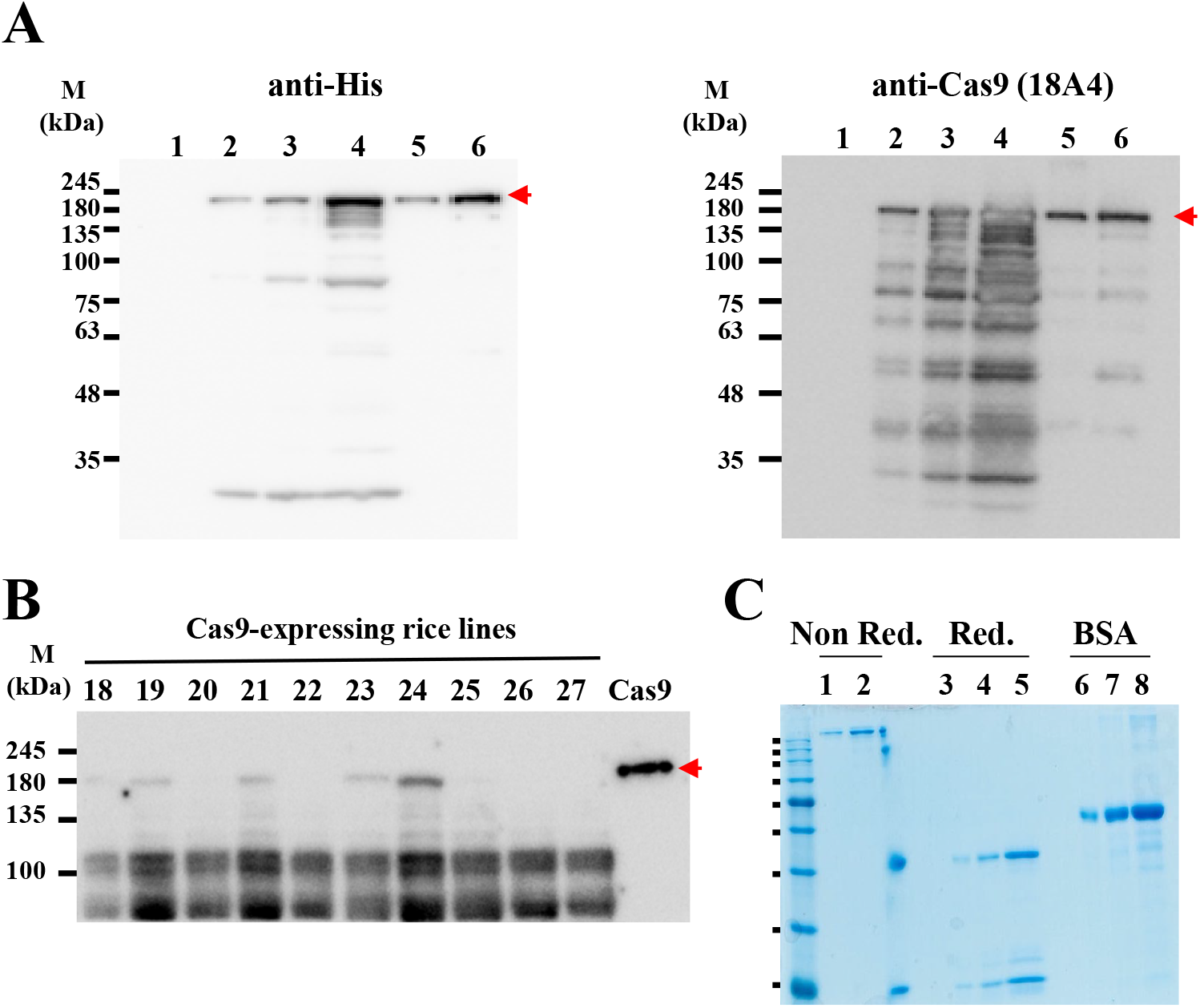
Western blotting for detection for spyCas9 expressed *E*.*coli* extract and IgG purification. (A) Detection of indicated amounts of spyCas9 expressed *E*.*coli* extract and purified SpyCas9 (in house or Toolgen SpyCas9 protein) using hybridoma culture media and anti-His. Lane 1; control Hela cell lysate, Lanes 2-4; SpyCas9 expressed *E*.*coli* extract (2.5, 4, and 10 μg), Lane 5; purified SpyCas9 (100ng), Lane 6; Toolgen SpyCas9 protein (100 ng). Exposed for 4 min and 30 seconds. (B) Detection of SpyCas9 protein overexpressed in rice (DJ strain) with hybridoma culture media. Lanes 15-27; Cas9 overexpressing rice (DJ) total protein (100ng), Lane 28; purified SpyCas9 (50ng). Exposed for one second. ***c***, SDS-PAGE results for purity determination of IgG purified SpyCas9 antibody– lanes 1-2, reduced conditions; lanes 3-5. non-reduced conditions; lanes 6-8, BSA for a loading control. Red arrows indicate SpyCas9 protein size. Red numbers indicate positive detection by anti-SpyCas9 antibody.

We also applied the supernatant from the hybridoma cell 18A4 to detect SpyCas that had been overexpressed in rice (*Dongjin*) after SpyCas9 DNA-mediated transformation. To examine whether the hybridoma supernatant detected codon-optimized SpyCas9 protein, the supernatant was used to identify high SpyCas9 expression lines out of SpyCas9 transgenic lines. The supernatant from the hybridoma cell 18A4 detected 6 lines, #16, 18, 19, 21, 23, and 24 out of 13 SpyCas9 overexpression lines (Fig. 3B).

### Anti-SpyCas9 antibody IgG purification from hybridoma cell 18A4 supernatant

The supernatant from the hybridoma cell 18A4 was tested to detect SpyCas9 (Figs. 3A-B). For specific binding to SpyCas9, anti-SpyCas9 IgG antibody was purified from the culture supernatant of hybridoma 18A4 via IgG purification. Purified anti-SpyCas9 antibody was separated on SDS-PAGE gels in nonreducing and reducing conditions (Fig. 3C). In the nonreducing condition, a band of a single size of over 245 kDa was accepted as anti-SpyCas9 antibody with an intact structure; in the reducing condition, two smaller fragments were obtained of the heavy and light chain of the antibody. Increasing the load of anti-SpyCas9 in gels in both nonreducing and reducing conditions, also increased the intensities of the observed bands for the intact antibody and reduced antibody components (Fig. 3C).

### Examination of spyCas9 protein with six other Cas9 proteins by ant-SpyCAs9 antibody

We examined whether the anti-SpyCas9 IgG antibody detected SpyCas9 protein expressed in *E. coli* and HEK293T cells. HEK293T cells were transfected with a SpyCas9 cloned into the PX459 vector. Two different dilutions (1:3,000 and 1:10,000) of anti-SpyCas9 IgG antibody successfully detected SpyCas9 proteins in HEK293T cell and Rosetta 2(DE3) pLysS *E. coli* cells as well as purified SpyCas9 protein (Fig. 4A). We also wanted to confirm whether purified SpyCas9 antibody could detect only SpyCas9 protein or any of the Cas9 variant proteins from other bacterial species. HEK293T cells were transfected with an additional seven mammalian expression vectors capable of expressing seven different Cas9 proteins: *Francisella tularensis* subsp. *novicida* Cas9; *Pasteurella multocida* Cas9; *Mycoplasma gallisepticum str. F* Cas9; *Nitratifractor salsuginis str*. DSM 16511 Cas9; *Campylobacter lari*. CF89-12 Cas9; *Parvibaculum lavamentivorans* Cas9; and *Streptococcus pyogenes* Cas9. The anti-SpCas9 antibody was specific for the SpyCas9 protein among these different Cas9 proteins, and the expression of all seven types of Cas9 protein was confirmed by a western blot using an HA-tag antibody (Fig. 4B).

**Fig. 4.**
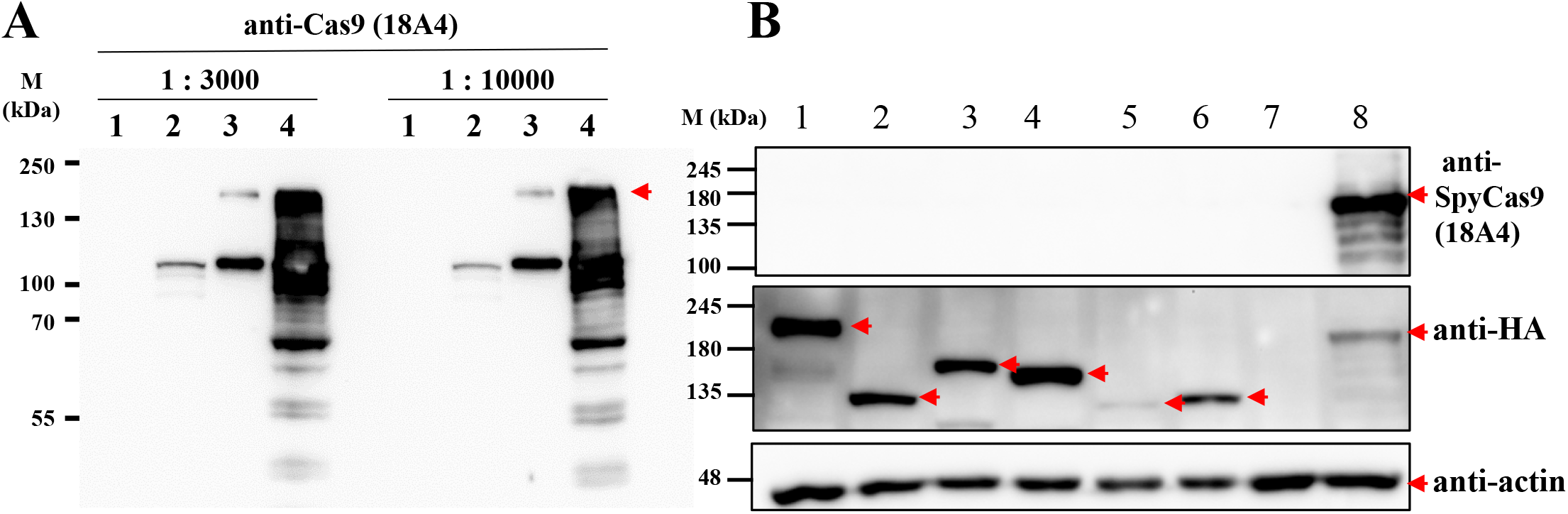
Western blotting; Detection for SpyCas9 expressed in HEK293T cell and Cas9 of six different species. (A) Anti-SpyCas9 antibody detected SpyCas9 protein. Lane 1; HEK293T lysates as a negative control, line 2; HEK293T transfected with SpyCas9 expression plasmid (50 μg total protein). Lanes 3; purified SpyCas9 (50 ng) and Lanes 4 SpyCas9 expressed in *Rosetta 2(DE3) pLysS* (50 μg total protein). The blotted membrane was probed by anti-SpyCas9 IgG antibody (18A4, 1:3000, 1:10000). Exposed for 6 minutes. (B) Detection of six different Cas9 proteins. HEK293T cells were transfected with each Cas9 expression plasmids. Lanes 1-6, six different Cas9 variant proteins; lane1, *Francisella tularensis subsp. novicida* Cas9; lane 2, *Pasteurella multocida* Cas9; lane 3, *Mycoplasma gallisepticum* str. F Cas9; lane 4, *Nitratifractor salsuginis* str. DSM 16511 Cas9; lane 5, *Campylobacter lari*. CF89-12 Cas9; lane 6, *Parvibaculum lavamentivorans* Cas9; lane 7, HEK293T lysates as a negative control; lane 8, SpyCas9 protein as a positive control. The blotted membrane was probed by anti-HA, anti-β-actin and anti-SpyCas9 IgG antibody (18A4, 1:3000). Red arrows indicate SpyCas9 protein bands.

The stability of the SpyCas9 protein was quantified in the cells after introduction of SpyCas9 protein into HEK293T cells by electroporation. The reduction in SpyCas9 abundance in cells was measured by harvesting cultures at 6, 18, 24, 30, 42, and 48 hours after transfection; harvested HEK293T cells were then analyzed by western blotting. Actin expression gradually increased with incubation time and proliferation of HEK293T cells (Fig. 5). In contrast, the abundance of SpyCas9 protein introduced into HEK293T cells decreased with incubation time. After 48 hours, SpyCas9 protein was reduced up to 30% compared to a 6-hour incubation (Fig. 5).

**Fig. 5.**
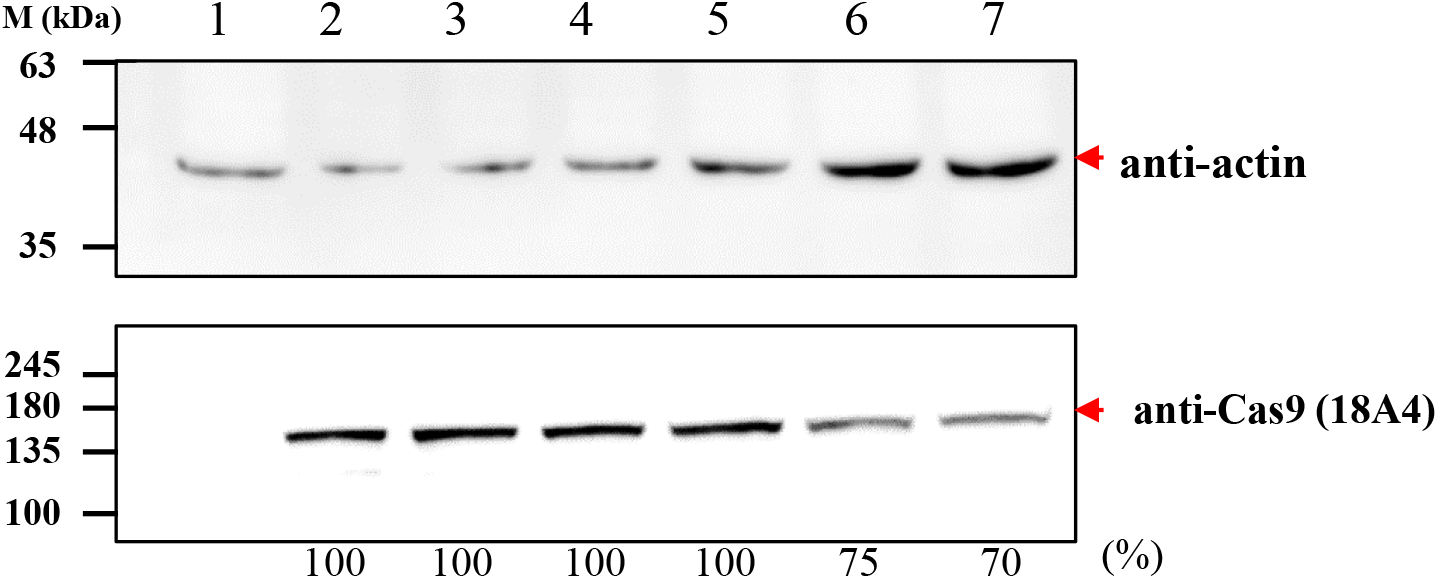
The stability of SpyCas9 protein after electroporation in HEK293T cells. Lane 1; HEK293T lysates as a negative control; lanes 2 to 7, HEK293T cells all were harvested from each well at 6, 18, 24, 30, 42, and 48 hours, respectively. Anti-actin reflected total crude protein amount of used HEK293T cells in an upper panel. All transfected SpyCas9 proteins from whole HEK293T lysates were detected by anti-SpyCas9 IgG antibody (18A4, 1:3000) in a bottom panel. Exposed for 20 and 60 seconds.

### Sequencing of three complementarity-determining regions (CDR)of heavy chain and light chain of anti-SpyCas9 (18A4)

The three CDR sequences of the variable region play an important role when an antibody recognizes an antigen (Supplementary Fig. S1). The hybridoma cell 18A4 was used for sequencing of three CDRs. The DNA sequence from the variant region to the CDR was amplified, and the amplicons were cloned and sequenced (Fig. 6) to identify the amino acid sequences of the three CDR and four FR regions of the variable region of antibody (Table 1).

**Table 1.**
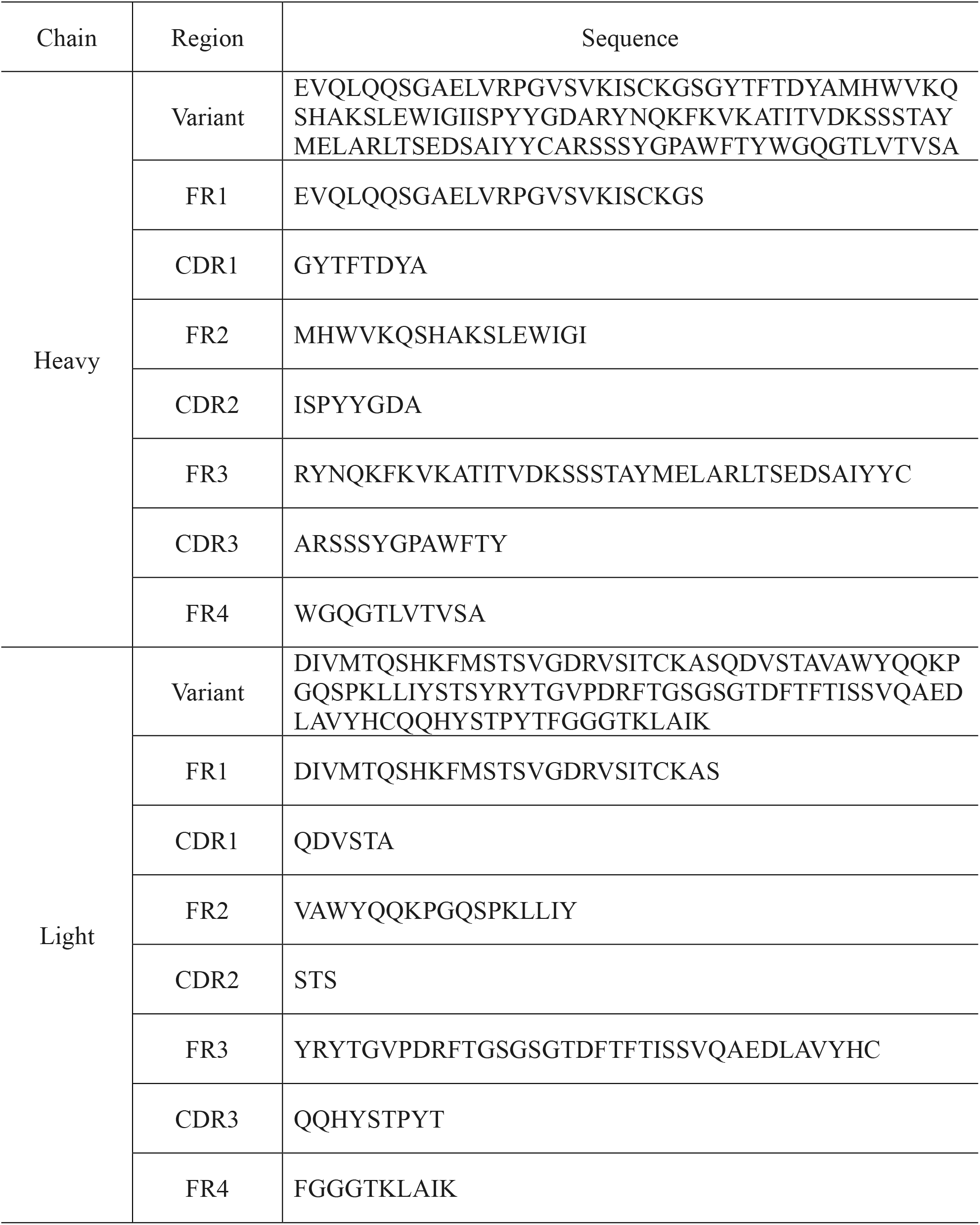
Sequence of VH and VL for SpyCas9 antibody.

| Chain | Region | Sequence |
| --- | --- | --- |
| Heavy | Variant | EVQLQQSGAELVRPGVSVKISCKGSGYTFTDYAMHWVKQ<br>SHAKSLEWIGIISPYYGDARYNQKFKVKATITVDKSSSTAY<br>MELARLTSEDSAIYYCARSSSYGPAWFTYWGQGTTLTVSA |
|  | FR1 | EVQLQQSGAELVRPGVSVKISCKGS |
|  | CDR1 | GYTFTDYA |
|  | FR2 | MHWVKQSHAKSLEWIGI |
|  | CDR2 | ISPYYGDA |
|  | FR3 | RYNQKFKVKATITVDKSSSTAYMELARLTSEDSAIYYC |
|  | CDR3 | ARSSSYGPAWFTY |
|  | FR4 | WGQGTTLTVSA |
| Light | Variant | DIVMTQSHKFMSTSVGDRVSITCKASQDVSTAVAWYQQKP<br>GQSPKLLIYSTSYRYTGVPDRFTGSGSGTDFTFTISSVQAED<br>LAVYHCQQHYSTPYTFGGGTKLAIK |
|  | FR1 | DIVMTQSHKFMSTSVGDRVSITCKAS |
|  | CDR1 | QDVSTA |
|  | FR2 | VAWYQQKPGQSPKLLIY |
|  | CDR2 | STS |
|  | FR3 | YRYTGVPDRFTGSGSGTDFTFTISSVQAEDLAVYHC |
|  | CDR3 | QQHYSTPYT |
|  | FR4 | FGGGTKLAIK |

**Fig. 6.**
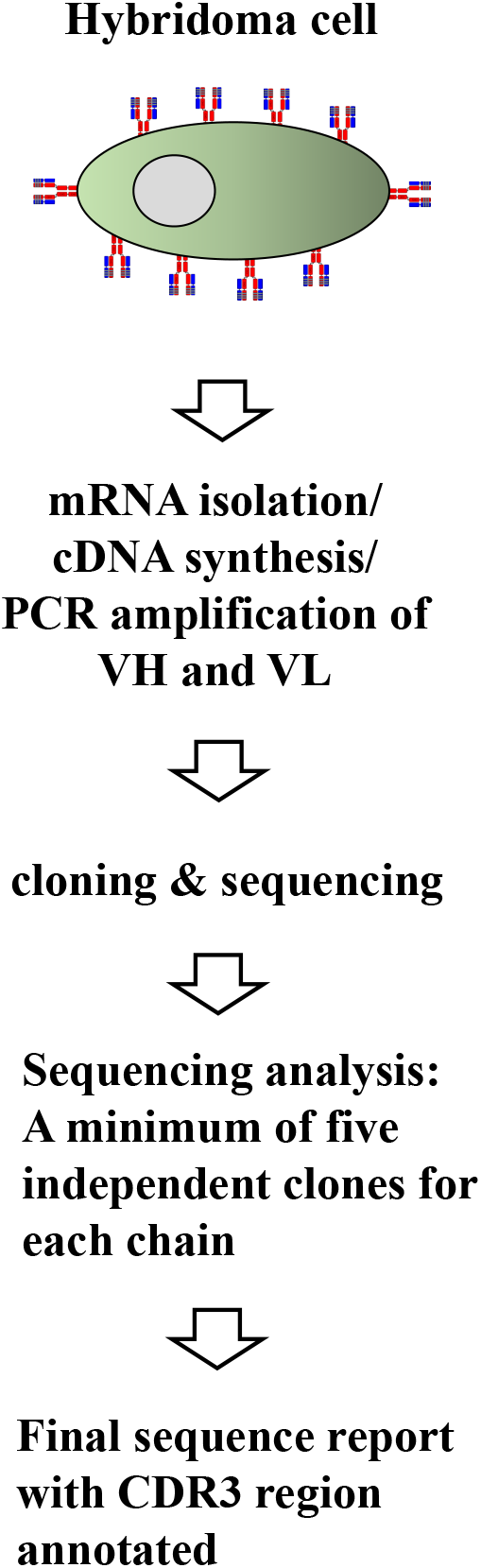
SpyCas9 antibody sequencing process. Complementary DNA was synthesized by using mRNA obtained from hybridoma 18A4 clone. Variable regions of heavy chain (VH) and light chain (VL) were amplified and cloned into pGEM®-T Vector. The colonies containing the variable regions of the heavy chain and the light chain were selected. For heavy chain and light chain, three CDR region sequences by sequencing at least over five clones were obtained.

## Discussion

Residual SpyCas9 in human cells needs to be quickly degraded following the precision cleavage at the targeted site and correct time, as the residual protein could result in cleavage of non-target sites as off-targeting effects (Hsu et al., 2013; Pattanayak et al., 2013). For example, increasing Cas9 protein levels by 2.6 fold led to a 2.6 fold increase in the number of off-target binding peaks in the genome (Wu et al., 2014). CRISPR technologies including SpyCas9, BE, and PE that are applied to gene or base editing in humans, animals, and plants, all require remove of residual protein to avoid such off-target effects. Therefore, we developed a qualified and sensitive anti-SpyCas9 antibody to be able to detect remaining SpyCas9 in cells. The high-sensitivity of the anti-SpyCas9 antibody helps to set a safety guideline for SpyCas9 usage in human therapies.

The anti-SpyCas9 antibody could also be employed to develop virus diagnosis kits based on the binding affinity between anti-SpyCas9 antibody and recombinant SpyCas9 protein. The principle of antigen-antibody binding has been widely adapted in multiple methods to rapidly diagnose diseases (Koczula and Gallotta, 2016); for instance, the ELISA system can demonstrate color- or fluorescence-based in 15 minutes (Wheeler, 2006) to enable a rapid diagnosis. In addition, many experimental techniques use antibodies as a tool for functional study, such as western blot, immunoprecipitation, and immunohistochemical staining. Therefore, it is important to produce specific monoclonal antibodies against versatile target proteins that can broaden the usage of these techniques. Cas9 proteins are versatile proteins that are applicable to many fields such as genome editing, transcriptional regulation, and therapeutic applications. Increased usage of SpyCas9 may require increased usage of detection systems based on the anti-SpyCas9 antibody described in this study.

## Author contributions

S.C designed, M.J.P and S.P conducted experiments, and J.P and S.C wrote the manuscript.

## Figure legends

**Fig. 1. Purification of SpyCas9 and immunization scheme. (A)** lane 1, 1 μg bovine serum albumin (BSA); lane 2, 2 μg BSA; lane 3, 1 μg SpyCas9; lane 4, 2 μg SpyCas9. The gel was stained with Coomassie Brilliant Blue. Recombinant SpyCas9 has an expected molecular mass of 166 kDa. (B) Immunization scheme for production of anti-SpyCas9.

**Fig. 2. Western blot of quantitative and specificity estimation for SpyCas9 antigen**. (A) SpyCas9 (12.5, 25, 50, 100, and 200 ng) was probed with hybridoma supernatants of selected cells (24H5, 9G8, 5G11, and 18A4). (B) Hela cell lysate (0, 5, 30, and 100 μg in lanes 1, 2, 3, and 4, respectively) was assessed for immunoreactivity against supernatants of selected hybridoma cell cultures (24H5, 9G8, 5G11, and 18A4).

**Fig. 3. Western blot of spyCas9 expressed in *E. coli* extracts and anti-SpyCas9 following IgG purification**. (A) Western blot of *E. coli* extract containing recombinant SpyCas9 (lanes 2, 3, and 4) and commercial SpyCas9 (lanes 5 and 6). (B) Western blot. (C) SDS-PAGE gel of IgG purified SpyCas9 in nonreducing conditions (0.1 and 0.4 μg in lane 1 and 2, respectively), and reducing conditions (0.1, 0.4, and 1.6 μg in lanes 1, 2, and 3 respectively).

**Fig. 4. Western blot of spyCas9 and six Cas9 variants in expressed HEK293T**. (A) western blot using anti-SpyCas9: Lane 1, *Francisella tularensis* subsp. *novicida* Cas9; lane 2, *Pasteurella multocida* Cas9; lane 3, *Mycoplasma gallisepticum str. F* Cas9; lane 4, *Nitratifractor salsuginis str*. DSM 16511 Cas9; lane 5, *Campylobacter lari*. CF89-12 Cas9; lane 6, *Parvibaculum lavamentivorans* Cas9; lane 8, *Streptococcus pyogenes* Cas9. (B) Western blot using an HA-tag antibody, lanes as above; the level of beta actin was checked to confirm that the same amount of sample was loaded.

**Fig. 5. The stability of SpyCas9 protein after electroporation in HEK293T cells**. Lane 1, HEK293T lysates as a negative control; lanes 2 to 7, HEK293T cells harvested at 6, 18, 24, 30, 42, and 48 hours, respectively. Transfected SpyCas9 proteins from whole HEK293T lysates were detected by anti-SpyCas9 IgG antibody (18A4, 1:3000) (upper panel). Anti-actin antibody was used to indicate the total amount of crude protein from transfected HEK293T cells (bottom panel).

**Fig. 6. SpyCas9 antibody sequencing process**. Complementary DNA was synthesized by using mRNA obtained from hybridoma 18A4 clone. Variable regions of heavy chain (VH) and light chain (VL) were amplified and cloned into pGEM®-T Vector. The colonies containing the variable regions of the heavy chain and the light chain were selected. For heavy chain and light chain, three CDR region sequences by sequencing at least over five clones were obtained.

---

**Fig. S1.**
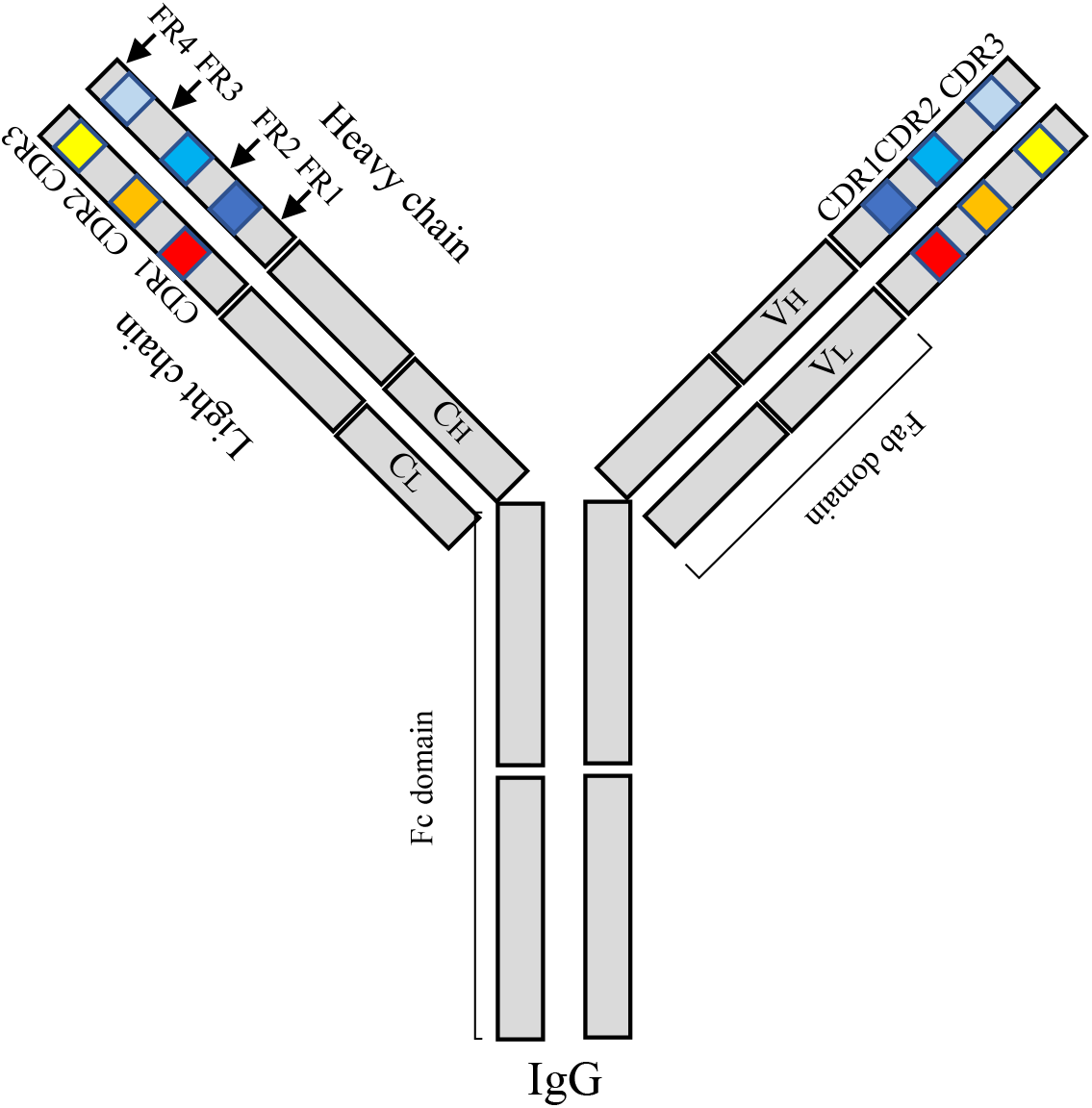
A schematic diagram of IgG.

